# Evaluating Variants of Uncertain Significance in Adult Knock-in Zebrafish: A Proof of Concept with a *COL1A2* Variant

**DOI:** 10.1101/2025.02.11.637642

**Authors:** Michiel Vanhooydonck, Sophie Debaenst, Eva Vanbelleghem, Hanna De Saffel, Delfien Syx, Patrick Sips, Paul J. Coucke, Andy Willaert, Bert Callewaert

## Abstract

Genomic variants of uncertain significance (VUS) impede clinical decision-making. In this study, we use a knock-in strategy in zebrafish to evaluate the *COL1A2* c.2123G>A VUS, identified in a 59-year-old female with recurrent fractures. Using prime editing, we obtained different zebrafish lines respectively harboring the VUS, a known pathogenic variant, or a known benign variant. Comprehensive skeletal phenotyping revealed no significant abnormalities in the zebrafish modeling the benign variant and the VUS, while zebrafish modeling the pathogenic variant showed scoliosis of the vertebral column, vertebral fusions, vertebral compressions, fractures, and increased mineralization of the notochord and intervertebral ligament. Our findings demonstrate for the first time, that *COL1A2* variant modeling in zebrafish models informs functional validation and shows potential for elucidating associated pathogenic mechanisms. This approach can be extended to study VUS in other genes.

## INTRODUCTION

Next- and third generation sequencing technologies greatly advanced the mapping of variation in the human genome. To assess whether specific variants are linked to disease, the American College of Medical Genetics (ACMG) criteria established guidelines, relying on evidence from population studies, computational data, functional testing, and segregation patterns to classify variants into five categories: pathogenic (class 5), likely pathogenic (class 4), uncertain significance (class 3), likely benign (class 2) and benign (class 1)^1^. Despite the progress in variant classification based on *in silico* prediction tools such as REVEL or CADD and population databases such as GnomAD and ClinVar, VUS remain the largest category of reported variants by diagnostic laboratories and are disclosed in up to 41% of reports concerning multigene panel tests^2^. Clinical interpretation of such variants is difficult, as extensive segregation analysis is not always possible. As a result, functional experimental data is required to reclassify these variants and determine the clinical relevance. *In vitro* analysis may be informative of a functional effect of the variant but may fail to establish a definite association between the variant and the disease. Therefore, proof of pathogenicity may be provided by modeling the variants in small laboratory animals.

Pathogenic variants in *COL1A2* are associated with either osteogenesis imperfecta (OI) or Ehlers-Danlos syndrome (EDS), two distinct connective tissue disorders. OI is a rare disorder characterized by increased bone fragility and skeletal deformities resulting from defects in collagen type 1 due to pathogenic variants in the *COL1A1* or *COL1A2* genes or genes involved in the processing of collagen type 1 chains. Specifically for *COL1A2*, missense variants, particularly glycine substitutions, disrupt the formation of the collagen type I triple helix, leading to an OI phenotype^3^, while most truncating mutations in *COL1A2* do not result in a phenotype due to nonsense-mediated mRNA decay (NMD)^4^. The cardiac valvular type of EDS is characterized by joint hypermobility, skin hyperextensibility and soft tissue fragility and caused by biallelic variants in *COL1A2* resulting in the absence of pro-α2(I) collagen chains^5^. Furthermore, the arthrochalasia type of EDS is associated with heterozygous exon 6 deletions (partial or complete)^6^ and more recently *COL1A2* variants have been identified in phenotypes with combined characteristics of both OI and EDS, so-called *COL1*-related overlap disorders^7^.

Zebrafish are widely used to investigate the pathogenic mechanisms of various diseases and have been used to test VUS^8–10^, usually based on early phenotypic readouts following injection of mutant mRNA containing the VUS into zebrafish with a gene knockout^11^. However, such approaches are complicated by a lack of dosage control and spatiotemporal regulation that modulates mRNA expression and precludes the investigation of adult phenotypes.

In this study, we provide the proof of principle using adult knock in zebrafish for reclassifying a VUS in the *COL1A2* gene.

## RESULTS

### Establishment of col1a2 VUS zebrafish model

The *COL1A2* c.2123G>A variant (NM_000089.4, NP_000080.2; p.(Arg708Gln)) has been identified in several patients with atypical femoral fractures^12^. Based on the ACMG classification (2021 criteria: BS2, BP6 and PP3)^1^, this variant has been classified as a VUS. To evaluate this variant, we modeled this VUS, a known benign (NM_000089.4:c.948C>T, NP_000080.2:p.(Gly316=)), and a known pathogenic variant (NM_000089.4:c.2738G>A, NP_000080.2:p.(Gly913Asp)) in zebrafish.

All three variants are conserved in zebrafish at the protein level and correspond to the *col1a2* c.918A>T/p.(Gly306=) (negative control), c.2093G>A/p.(Arg698Gln) (VUS) and c.2645 G>A/p.(Gly882Asp) variants (negative control), respectively (GRCz11, **Supplementary Figure** , **Supplementary Table**. To enhance readability, we refer to negative control as *col1a2*^*SNP/+*^, the VUS as *col1a2*^*VUS/+*^ and the positive control as *col1a2*^*mh15/+*^. The VUS and the benign control were introduced in the zebrafish genome using prime editing^13^, while the known pathogenic variant was previously established and characterized in our lab as *col1a2*^*mh15/+*^ zebrafish mutants^14,15^.

### Skeletal phenotyping reveals no abnormalities in the VUS zebrafish model

We used Alizarin Red S mineral to visualize bone tissue in the VUS (n=20), negative control (n=13) and positive control (n=11) zebrafish lines. *Col1a2*^*mh15/+*^ mutant zebrafish displayed several skeletal abnormalities, including an increased sagittal curvature of the spine (mimicking scoliosis), more vertebral fusions (marked by reduced intervertebral space) and compressions (marked by narrower vertebral bodies), and more vertebral fractures (marked by fracture lines and/or callus formation). Furthermore, *col1a2*^*mh15/+*^ mutants show increased mineralization of the notochord and intervertebral ligament (IVL) following staining for mineral deposits^14,15^ as well as an increased eye diameter, which correlated with standard length^16^. In contrast, the *col1a2*^*VUS/+*^ zebrafish, showed no significant skeletal differences comparable to the *col1a2*^*SNP/+*^ negative controls (**Figure 1**).

**Figure:**
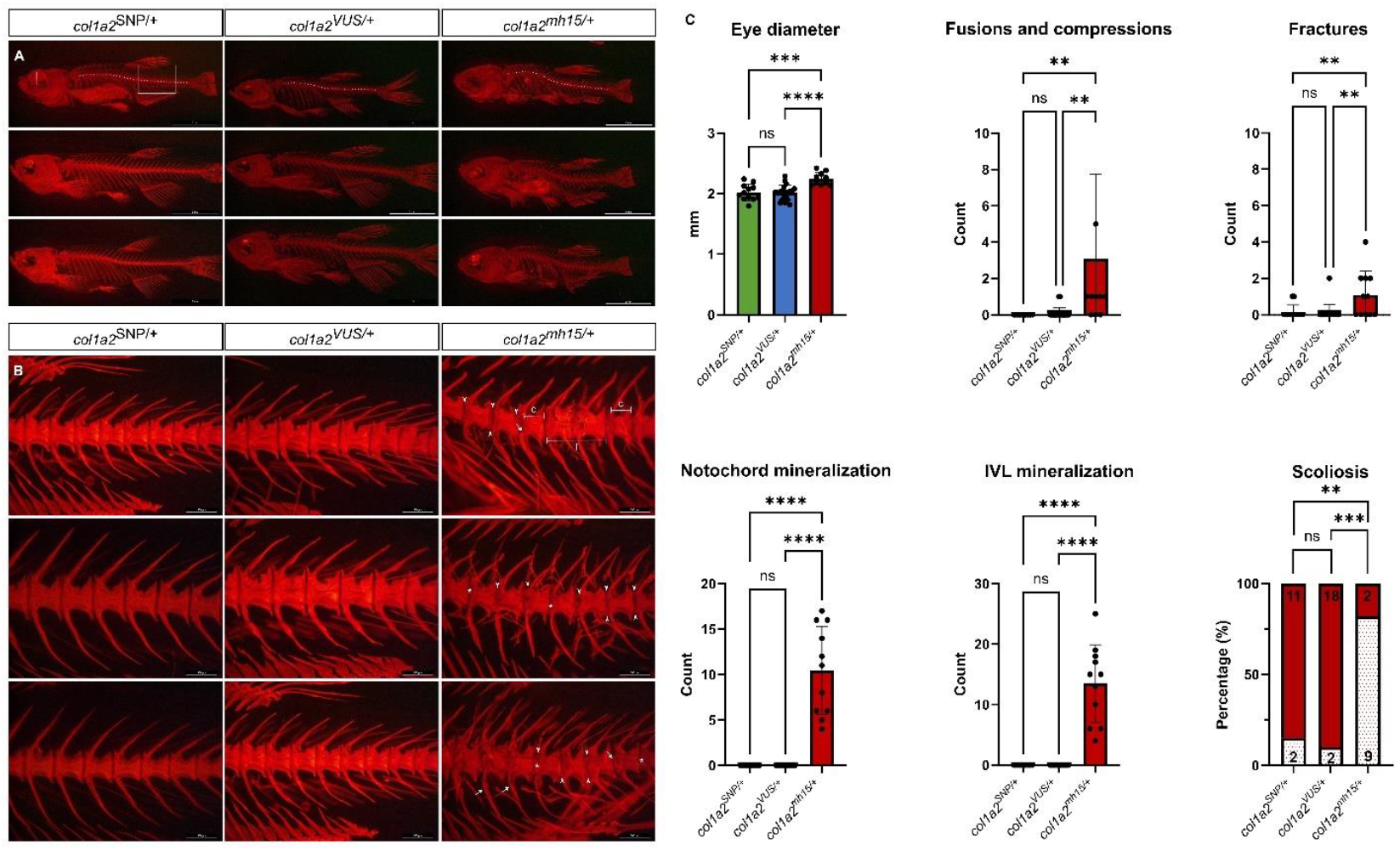
Evaluation of skeletal anomalies in *col1a2*^*SNP/+*^, *col1a2*^*VUS/+*^ and *col1a2*^*mh15/+*^ zebrafish. (A) Overview images of three representative zebrafish for each genotype. The white dotted line traces the curvature of the vertebral column, and the eye diameter is marked with a white line. The white box highlights the region magnified in panel (B). Scalebar 5 mm, 7.3x magnification **(B)** A detailed view of the caudal vertebrae shows no skeletal abnormalities in the *col1a2*^SNP/+^ and *col1a2*^VUS/+^ zebrafish while the positive control *col1a2*^mh15/+^ shows vertebral fusions (f), compressions (c), as well as mineralization of the notochord (*) and intervertebral ligament (IVL) (arrowhead). Scalebar = 500 μm, 80x magnification **(C)** Skeletal phenotyping of the zebrafish. The X-axes represent the genotype. In the graphics for eye diameter the Y-axis represents the absolute diameter. For fusions and compressions, fractures, notochord mineralization, and IVL the Y axis represents the total number of occurrences of each deformity in the vertebral bodies per individual, excluding the first four vertebrae (which form the Weberian apparatus) and the last four vertebrae (preural and ural vertebrae) from the analysis. For scoliosis, zebrafish exhibiting scoliosis are shown in white, while those without scoliosis are shown in red. The number of analyzed fish: *col1a2*^*SNP/+*^ = 13, *col1a2*^*VUS/+*^ = 20, *col1a2*^*mh15/+*^ = 11. Asterisks denote the level of significance: **p<0.01, ***p<0.001, ****<0.0001. Data are represented as the mean ± standard deviation.

## DISCUSSION

This study provides a proof of concept to evaluate the pathogenicity of VUS using a knock-in strategy in zebrafish. We evaluated the *COL1A2* c.2123G>A p.(Arg708Gln) variant, a VUS reported in individuals with atypical femoral fractures^12^. Zebrafish harboring the homologous variant are phenotypically not different from similar *col1a2*^*SNP/+*^ negative control zebrafish, while the positive control *col1a2*^*mh15*/+^ clearly shows skeletal defects, allowing us to classify the *COL1A2* c.2123G>A p.(Arg708Gln) variant as (likely) benign. Of note, when our study started the variant was classified as a variant of unknown significance, while an updated classification using the ACMG guidelines now categorizes the variant as ‘likely benign’ (BS2 and BP6 criteria)^1,17^, supporting our findings. Hence, our study allows to add the BS3 criterion to the variant classification.

For such studies, the use of appropriate controls is of utmost importance, taking into account the mutational mechanisms. We therefore opted to introduce a heterozygous pathogenic glycine substitution to obtain a positive control line. Missense variants, particularly glycine substitutions within the alpha-helical domain of the *COL1A2* gene, cause OI while a heterozygous null allele is unlikely to cause disease and biallelic null alleles typically result in a valvular phenotype, unrelated to OI^4^.

Zebrafish has already demonstrated its potential as a modeling organism to study the functional impact of VUS^9,18^. However, most approaches establish knockout models to evaluate the phenotypic consequences. While this provides valuable insights into the gene’s role, it does not directly test the specific patient-derived variant, potentially limiting the relevance of the findings^8^. Variant analysis has been performed by injecting mRNA containing the VUS into wild-type zebrafish embryos^9^ or zebrafish KO models for the gene of interest^11^, but has many limitations. *In vitro* transcription and purification of large mRNA molecules can be challenging. While plasmid-based approaches may address some of these issues, they present limitations such as lower protein expression, the need for nuclear localization, and the risk of insertional mutagenesis. Both mRNA and plasmid-based methods are also prone to toxicity effects, lack control over spatiotemporal expression, and are restricted to early development phenotypes due to transient expression, as mRNA transcripts or plasmids diminish with each cell division. In contrast, generating a knock-in (KI) line, coupled with well-validated positive and negative controls, offers a more robust strategy to accurately assess the pathogenicity of VUS.

Nevertheless, the major limitations associated with generating stable KI zebrafish lines is the extended timeline, requiring at least six to nine months, and low editing efficiencies, although recent advancements in genome editing technologies, such as prime-editing improved efficiency^13,19^. As an alternative to homology-directed repair (HDR), prime-editing further reduces the risk for off-target effects and reduces the risk of non-homologous end joining as it does not require a double-stranded break^19,20^.

Our findings demonstrate that *COL1A2* VUS testing in zebrafish is feasible for skeletal abnormalities and could be evenly successful for *COL1-*related overlap disorders. Hence, our established knock in generation pipeline holds promise to advance our understanding of the pathogenic mechanisms underlying OI, EDS or other *COL1*-related disorders. Furthermore, our proof-of-concept offers potential to investigate VUS in other genes offering valuable diagnostic insights, enabling OI subtype stratification, providing predictive value for disease progression, and informing treatment decisions.

## DATA AVAILABILITY STATEMENT

All data are available from the authors upon request.

## ACKNOWLEDGMENTS

The authors would like to thank the Zebrafish Facility Ghent (ZFG) Core at Ghent University, and particularly Karen Vermeulen for the diligent care for the zebrafish.

## ETHICS DECLARATION

The zebrafish (*Danio rerio*) AB line was maintained and handled in accordance with the Animal Welfare Legislation, EU Directive 2010/63/EU (European Commission, 2016). The study was approved by the local committee on the Ethics of Animal Experiments of Ghent University Hospital, Ghent, Belgium with permit number ECD20-41 and ECD23-27.

