## Supplementary Material for "Evaluating Variants of Uncertain Significance in Adult Knock-in Zebrafish: A Proof of Concept with a *COL1A2* Variant"

### SUPPLEMENTARY METHODS

#### *Zebrafish Husbandry and maintenance*

The zebrafish (*Danio rerio*) AB line was maintained and handled in accordance with the Animal Welfare Legislation, EU Directive 2010/63/EU (European Commission, 2016). The study was approved by the local committee on the Ethics of Animal Experiments of Ghent University Hospital, Ghent, Belgium with permit number ECD20-41 and ECD23-27.

#### *Generation of the zebrafish models*

The *col1a2*$^{SNP/+}$ and the *col1a2*$^{VUS/+}$ models were obtained using CRISPR/Cas9 gene editing technology following protocols described previously[1]. Utilized pegRNAs and genotyping primers are listed in **Supplementary Table**. The *col1a2*$^{mh15/+}$ zebrafish line was obtained from the lab of Matthew P. Harris[2]. Adult mutants were obtained through outcross breedings with AB zebrafish. Embryos were raised in an incubator at 28°C, after which they were transferred in equal densities to 3,5L tanks in the Zebrafish Core Facility Ghent (ZFG), where they were grown until adulthood.

#### *Genotyping*

At 3 months post-fertilization, adult zebrafish were fin-clipped for DNA extraction. Genomic DNA was isolated using a 50mM NaOH solution and subsequently neutralized with 1M Tris-HCl. Polymerase chain reaction (PCR) amplification was performed using specific primers, followed by enzymatic purification of the PCR products with exonuclease I and alkaline phosphatase. Purified PCR products were prepared for sequencing, which was performed using a standard Sanger sequencing protocol. The resulting sequences were further purified using bead-based cleaning methods to ensure high-quality data. Sequence analysis was conducted using FinchTV software. All genotyping primers and PCR protocols are listed in **Supplementary Table**. Following protocols were used: TD60-48 (2’ 94°C, 12 x (30’” 94°C, 30” 60*°C, 1’ 72°C), 25 x (40” 94°C, 40” 48°C, 30” 72°C), 10’ 72°C). TD62-50 (2’ 94°C, 12 x (30’” 94°C, 30” 62*°C, 1’ 72°C), 25 x (40” 94°C, 40” 50°C, 30” 72°C), 10’ 72°C), lowering the annealing temperature with one degree for the first twelve steps.

#### *Alizarin red S mineral staining*

Adult zebrafish were stained using the Alizarin Red S mineral staining protocol as described by Beck *et al.* (2021)[3]. Fish were euthanized with a 25× tricaine solution and subsequently transferred to a fixative solution (5% NBF, 5% Triton, 1% KOH). Specimens were incubated in the fixative for 10 days at room temperature. Following fixation, the fish were transferred to an enhancement solution (20% ethylene glycol, 5% Triton, 1% KOH) and incubated for 24 hours at room temperature. Specimen were immersed in a bone staining medium (20% ethylene glycol, 1% KOH) for 30 minutes before being transferred to the Alizarin Red S staining solution (20% ethylene glycol, 1% KOH, 0,05% Alizarin Red S). After staining, scales were gently removed from the fish using a cotton swab. The specimens were then immersed in a destaining solution (20% Tween 20, 1% KOH) for 24–72 hours to enhance clarity and remove excess dye. Fish were progressively transferred through a graded series of glycerol solutions and stored in 100% glycerol for long-term preservation.

Stained zebrafish specimens were imaged using a Leica M165FC binocular stereomicroscope equipped with LAS V4.3 software (Leica Microsystems). Lateral views were primarily utilized for skeletal assessments, while ventral views were employed for the evaluation of spinal curvature. Imaging was performed using both brightfield and fluorescent modes, with fluorescence enabled by the DSR DsRed Dichroic Mirror filter. This imaging approach allowed for a comprehensive examination of skeletal features and the identification of deformities. The number of fusions and compressions, and fractions occurring in the entire vertebral column were counted for each individual fish and plotted per fish for all three zebrafish lines. For notochord mineralization and IVL mineralization, the number of mineralized IVL or notochord sections were counted over the entire vertebral column and plotted per zebrafish for all three lines. For scoliosis, the zebrafish were positioned dorsally and if the vertebral column deviated from a straight line, the fish was assigned to the scoliosis group. For each zebrafish line, the number of zebrafish with and without scoliosis were counted and reported as a percentage.

#### *Statistical analysis*

Statistical analysis of fish with scoliosis was done using the Fisher’s exact test. All other parameters were evaluated using one-way ANOVA followed by Tukey’s multiple comparison test. All analysis were performed in Graphpad Prism 10.2.3.

### SUPPLEMENTARY FIGURE

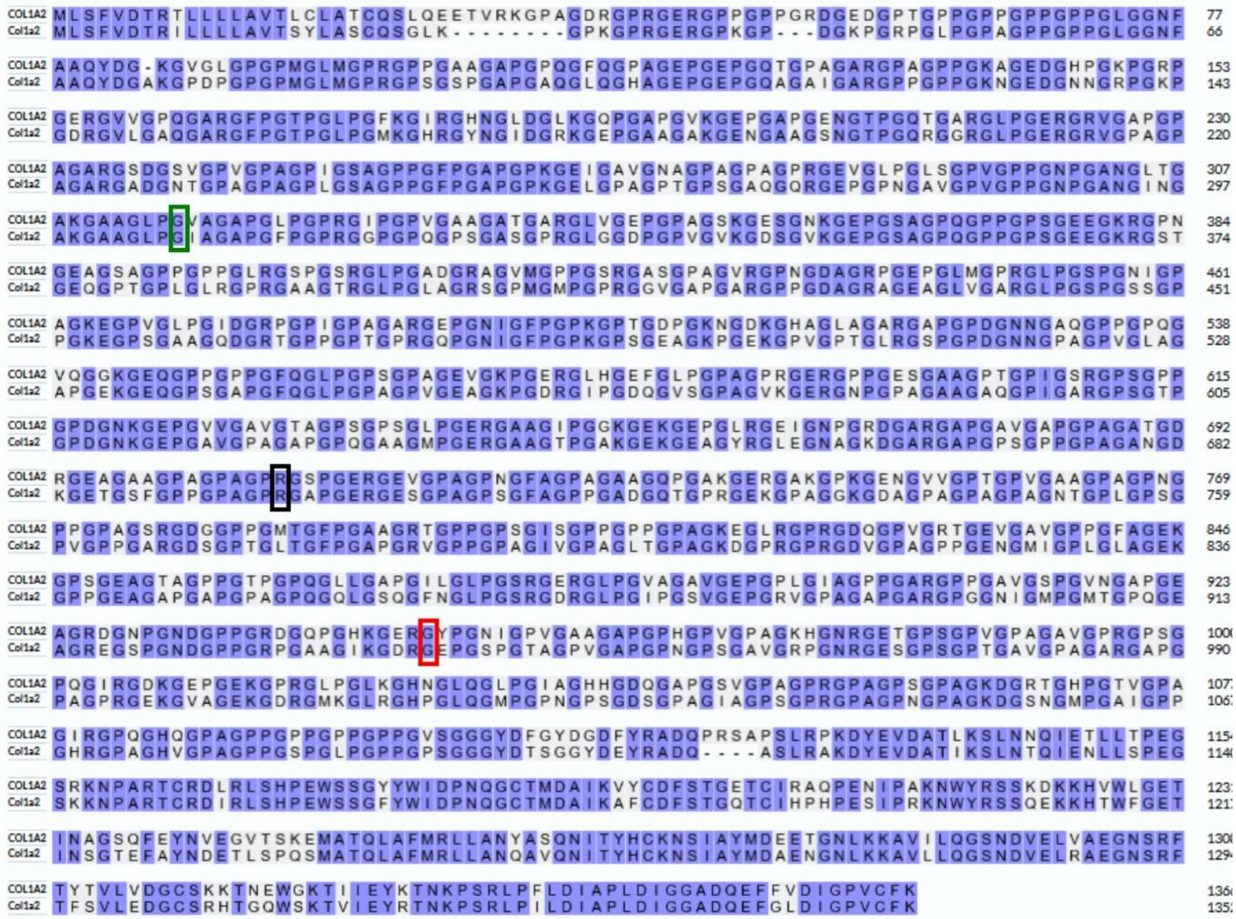


**Supplementary Figure**: Protein alignment of human *COL1A2* with zebrafish *Col1a2*. Conserved amino acids are indicated in purple. Conserved variants are indicated in green, black and red for the negative control, VUS and positive control respectively.

### SUPPLEMENTARY TABLES

**Supplementary Table:** Variant details of human and zebrafish, pegRNA sequences, genotyping primers, PCR protocol, and cDNA primers utilized for all targets. For pegRNAs, desired edits are colored red, spacer region is underlined, PBS is bold and RTT is blue.

| | TARGET GENE | HUMAN VARIANT | ZEBRAFISH VARIANT | pegRNA | GENOTYPING PRIMERS | PCR PROTOCOL |
|---|---|---|---|---|---|---|
| negative control | ***COL1A2*** | c.948C>T<br>p.Gly316= | c.918A>T<br>p.Gly306= | CCAGGGGCACCAGCAATTCCGTTTTAGAGCT<br>AGAAATAGCAAGTTAAAATAAGGCTAGTCCG<br>TTATCAACTTGAAAAAGTGGCACCGAGTCGG<br>TGCACAGGGTCTTCCTGGT**ATTGCTGGTG** | F: GGCTGTTAATTGTTTTCTTAGGCA<br>R: ACGTTGTGAATGCAGCAGTTAG | TD60-48 -1°C |
| VUS | ***COL1A2*** | c.2123G>A<br>p.Arg708Gln | c.2093G>A<br>p.Arg698Gln | AAACTCACAGGGGCACCACGGTTTTAGAGCT<br>AGAAATAGCAAGTTAAAATAAGGCTAGTCCG<br>TTATCAACTTGAAAAAGTGGCACCGAGTCGG<br>TGCTGCTGGTCCTCAT**GGTGCCCCTG** | F: ACTTGCAAGACTCTCAAAAGAACT<br>R: CATGGACTTACAGGGGGTCC | TD60-48 -1°C |
| positive control | ***COL1A2*** | c.2738G>A<br>p.Gly913Asp | c.2645 G>A<br>p.Gly882Asp | | F: AATAGGGAGCACCTGGACCT<br>R: ATTCCAGCAGCACCAGGAC | TD62-50 -1°C |
